# Time-Resolved Resonance Raman Spectroscopy of Retinal Proteins with Continuous-Wave Excitation. A Fundamental Methodology Revisited

**DOI:** 10.64898/2026.01.18.700169

**Authors:** Anna Lena Schäfer, Cristina Gellini, Rolf Diller, Katrina T. Forest, Uwe Kuhlmann, Peter Hildebrandt

**Affiliations:** Technische Universität Berlin, Institut für Chemie, Straße des 17. Juni 135, D-10623 Berlin, Germany; Università di Firenze, Dipartimento di Chimica “Ugo Schiff”, Via della Lastruccia 3-13, 50019, Sesto Fiorentino, Italy; RPTU Kaiserslautern, Fachbereich Physik, Erwin Schrödinger Str., Geb. 46, D-67663 Kaiserslautern, Germany; University of Wisconsin-Madison, Department of Bacteriology, 1550 Linden Dr., Madison, WI 53706, USA

**Keywords:** resonance Raman, time-resolved, confocal spectrometer, retinal protein, bacteriorhodopsin

## Abstract

Time-resolved resonance Raman spectroscopy with continuous-wave excitation is a fundamental technique that has contributed substantially to the understanding of structure and dynamics of bacteriorhodopsin and related retinal proteins. However, the underlying principles were developed about fifty years ago for instrumentation that is hardly in use any more. Thus, the adaptation of the technique to current state-of-the art equipment is needed to satisfy the increasing demand for the spectroscopic characterization of microbial retinal proteins. In this work, we focus on pump-probe time-resolved resonance Raman experiments with a confocal spectrometer using a rotating cell. We discuss the boundary conditions that fulfill the fresh sample conditions and the photochemical innocence of the probe beam as a prerequisite for studying parent or intermediate states of retinal proteins that undergo a cyclic photoinduced reaction sequence. For the measurements of intermediate states and reaction kinetics, pump-probe experiments are required in which the two laser beams hit the flowing sample with a defined but variable delay time. An appropriate set-up for such two-beam experiments with a confocal spectrometer is proposed and tested in time-resolved experiments of bacteriorhodopsin. The comparison with the results obtained with previous classical slit spectrometers with 90-degree-scattering illustrates the advantages and disadvantages of the confocal arrangement. It is shown that modern confocal spectrometers substantially decrease the spectra acquisition time but require a more demanding optical set-up. Furthermore, the extent of photoconversion by the pump beam is lower than for the 90-degree-scattering arrangement which lowers the accuracy of kinetic measurements.

## 1 Introduction

Resonance Raman (RR) spectroscopy is an important tool to study biological photoreceptors, since it allows probing structural and reaction kinetics over a wide dynamic range.^[1]^ The technique offers insight into structural details of the cofactors that are beyond the resolution of X-ray crystallography and even encompasses intermediate and instable states of the photoreceptors. Particularly appropriate research objects are bacterial retinal proteins that upon light absorption run through cyclic photoinduced reaction sequences which return to the initial state (Fig. 1A).^[2]^

**Figure 1.**
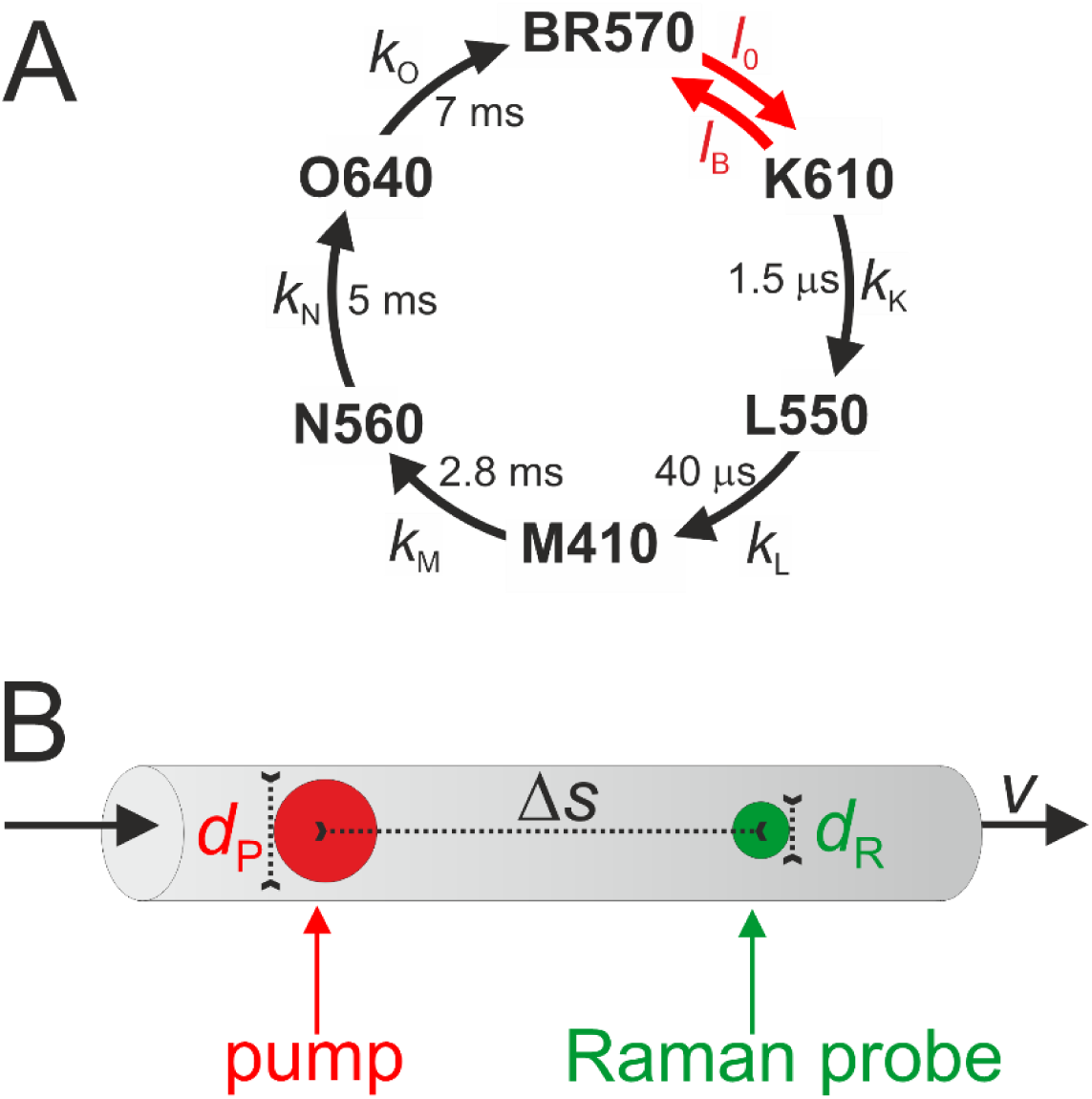
Basis of cw pump-probe TR RR experiments on BR. (**A**) Simplified scheme of the photocycle of BR. Spectral and kinetic data was taken from the literature.^[2,3]^ The individual states are indicated by the commonly used notation with the numbers indicating the absorption maxima. The photochemical and thermal reactions are characterized by the red and black arrows/symbols, respectively. The approximate life times of the individual reaction steps are indicated. **B**, Principle of TR pump-probe RR spectroscopy using a flowing sample with the velocity *v* and the pump and Raman probe beams separated by the distance Δ*s*. The beam diameter, defined by the intensity drop to 1/*e*^2^of the maximum laser power, must be larger for the pump (*d*_P_) than for the Raman probe beam (*d*_R_).

Here bacteriorhodopsin (BR) has played a prominent role in the past.^[2,4,5]^ Its discovery nearly coincided with the first attempts to apply RR spectroscopy to biological systems.^[1]^ Soon it turned out that RR spectroscopy could provide valuable information about the chromophore structure as well as the reaction dynamics and mechanism of BR.^[6–10]^ But more than that, due to its high stability in its native two-dimensional crystalline lattice (purple membrane), availability in large amounts, and the convenient kinetic properties, BR served as an ideal model system to develop and to test key methodological approaches in RR spectroscopy and also in many other spectroscopic techniques.^[11–17]^

Based on BR, the concept of time-resolved (TR) pump-probe RR spectroscopy with continuous-wave (cw) lasers was elaborated in the seminal works of the Stockburger and Mathies groups in the late seventies and early eighties.^[6,7,18–32]^ In these approaches, the sample flows through the pump and probe laser beams, separated by a well-defined distance Δ*s* (Fig. 1B). Velocity of the sample flow *v* and photon fluxes of the two beams define the two essential criteria that must be fulfilled: (i) repetitive measurement events must always start with the non-photolyzed sample (fresh sample condition) and (ii) the Raman probe beam must induce photoconversion only to a negligible extent (photochemical innocence of the probe beam).

In general, TR RR spectroscopy with cw laser excitation is restricted to a time-resolution of ca. 1 μs and thus complementary to TR RR spectroscopy using pulsed lasers that operate in the nano- and picosecond time range,^[1,33,34]^ and stimulated Raman spectroscopy that allows probing events even in the femtosecond time range.^[17,35]^ However, the advantage of cw laser measurements is the distinctly lower photon flux compared to pulsed laser excitation, which substantially reduces the risk of photodamage of the sample. Furthermore, cw RR spectroscopy allows for a high spectral resolution and yields a better signal-to-noise ratio than experiments with pulsed laser excitation when comparing the same accumulation time.

Whereas for a long time the studies of bacterial retinal proteins were largely restricted to BR, about 25 years ago the number of newly discovered microbial rhodopsins increased enormously and with it the demand for spectroscopic characterization.^[2,36]^ Thus, the field is currently witnessing a renaissance in TR RR spectroscopy of retinal proteins.^[37–48]^ However, not in every case are the critical principles of TR RR spectroscopy adequately considered. This is partly due to the progress in instrumentation, which has revolutionized Raman spectroscopy.^[49,50]^ The slit spectrometers that were available forty years ago facilitated the proper two-beam alignment and thus the fulfillment of the two abovementioned criteria of TR RR spectroscopy. This spectrometer generation has meanwhile largely been replaced by confocal spectrometers which substantially improve the collection of the scattered light and reduce the time for spectra acquisition enormously. Disadvantages of confocal setups, however, are the high photon flux in the focal spot of the probe beam and the severe difficulties in alignment of a two-beam experiment (vide infra).

The present work is directed to optimize the approach of a cw pump-probe TR RR experiment with a confocal detection system. Possibilities and limitations for the application to retinal proteins will be demonstrated and discussed in comparison with previous results obtained with classical slit spectrometers. The paper is organized as follows. First, we will recall the critical criteria of pump-probe TR RR spectroscopy, define the range of experimental parameters, such as probe beam laser power and velocity of the sample flow, and describe how to meet these criteria in comparison with those for slit spectrometers. Second, we propose a practical solution for a two-beam set-up using confocal spectrometers and discuss difficulties of the alignment. The subsequent section is dedicated to the application to the benchmark photoreceptor BR (Fig. 1A). We compare results obtained with confocal and slit spectrometer and, finally, discuss advantages and disadvantages of the spectroscopic setups.

## 2 Materials and Methods

Most RR spectra were recorded using a confocal Raman spectrometer (Horiba, LabRam HR-800) equipped with a liquid-nitrogen cooled CCD detector (Horiba, LN-BIUV-2048). The spectral resolution was 1 – 2 cm^−1^ and the wavenumber increment per pixel ca. 0.2 cm^−1^. The probe laser line was focused onto the sample to a spot of 4 μm diameter using a microscope objective with a numerical aperture of 0.35. Spectra were calibrated to an accuracy of ca. 0.5 cm^−1^ using the Raman bands of toluene as a reference.

RR spectroscopy based on conventional spectrometers was carried out with two different devices. As a scanning system, a Spex 14018 double monochromator, equipped with EMI 9659 QB photomultiplier, and a photon counting system (Ortec) were used. Overview spectra in the region between 700 and 1700 cm^−1^ were recorded with a step width of 1 cm^−1^. For the narrow C=C stretching region a step width of 0.2 cm^−1^ was used. The dwell time at each step was 1 s. In all cases the spectral bandwidth was ca. 4 cm^−1^.^[28]^ Alternatively, a U1000 spectrometer, operated as a spectrograph and equipped with a liquid-nitrogen-cooled CCD detector (Instruments SA), was employed. The spectral slit width was 2.8 cm^−1^.^[51]^

Details of the pump-probe experiments are discussed in the Results and Discussion section, and, in the case of slit-spectrometer set-ups, were reported in earlier works.^[28,31]^ For all experiments, Kr and Ar ion cw lasers (Coherent, CR and Innova series) with wavelengths at 458, 488, 514, 530, 568, and 647 nm were used for pumping and probing. Measurements were conducted at ambient temperature. The samples were filled into a rotating quartz cuvette. Data processing, such as baseline corrections, were conducted using the OPUS 7.8 (Bruker) software and in-house programs. No smoothing procedures were applied to the experimental spectra. Band fitting using Lorentzian band-shapes and component analysis was conducted using in-house programs.^[52,53]^

Purple membrane (BR) and halorhodopsin (HR) was prepared as described elsewhere.^[54,55]^ For RR experiments with the confocal and slit spectrometer set-ups we used concentrations corresponding to an optical density at the absorption maximum of ca. 10 and 1.0, respectively. The solution was buffered to pH 7.0 or 8.0 by Tris buffer.

## 3 Results and Discussion

The present study employs a rotating cell that was introduced into Raman spectroscopy already fifty years ago (Figs. 2, 3).^[56]^

**Figure 2.**
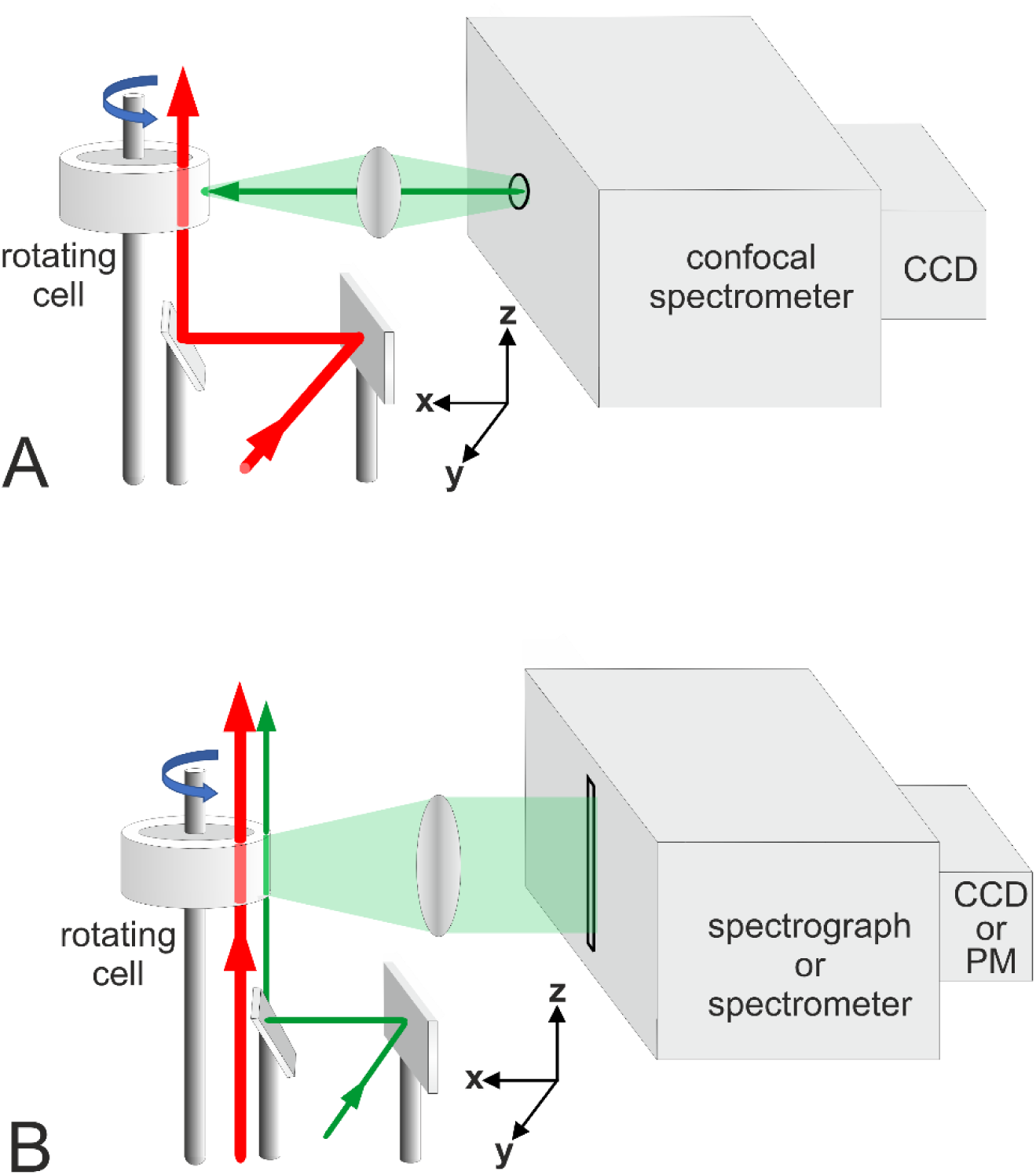
Sketch of the setups of TR RR pump-probe experiments with a (**A**) confocal spectrometer and (**B**) a slit spectrometer. The sample solution is filled into the rotating cell.

Upon rotation a solution layer is formed at the inner cell wall which the two spatially displaced laser beams passed through. By controlling the rotation speed of the cuvette and the timing between pump and probe pulses, this device permits the user to define the time delay δ between the pump and probe beams reaching the sample. Notably, it requires only small amounts of sample. In contrast, the capillary flow system, which has also been used in TR RR experiments previously, can hardly be employed in confocal setup since the high concentrations that are required (*vide infra*) limit the velocity of the sample flow substantially and are associated with large amounts of sample.

### 3.1. General principles of cw TR RR spectroscopy

We first consider a single-beam probe-only experiment with a retinal protein that runs through a reversible photocycle. The objective of the probe-only experiment is to measure the RR spectrum of the parent (dark) state which requires (i) a low photon flux of the probe beam such that the probability of photoconversion is minimal, and (ii) regeneration of the irradiated sample volume before it enters the laser beam again.^[1,6,25,31]^ Numerically, photoconversion is given by the photochemical rate constant *l*0 (in s^−1^), according to

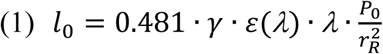

as described in the Supporting Information.^[29,57,58]^ Here γ, *λ*, and ε(*λ*) denote the photochemical quantum yield, the wavelength of the Raman probe beam (in nm), and the extinction coefficient of the retinal chromophore at *λ* (in *L* · *mol*^−1^ · *cm*^−1^), respectively. For 514.5 nm excitation and exemplary values for γ = 0.5 and ε(514) = 30000 *L* · *mol*^−1^ · *cm*^−1^),^[57]^ Eq. (1) yields

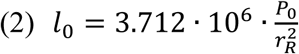

Then the main parameters to control *l*_0_are the laser power *P*_0_ (in mW) and the radius of the Raman probe beam *r*_*R*_ (in μm). The quantity *r*_*R*_ is only a few μm in confocal set-up but tens of μm in slit spectrometer set-ups, such that for the same laser power the photon flux is 100 to 1000 times higher in confocal setups. Thus, in confocal measurements, *P*_0_ must be substantially reduced to avoid formation of unwanted intermediates in the probe beam (Supporting Information, Fig. S1, Table S1). We may use the photoconversion parameter as a convenient quantity to quickly estimate the extent of photoconversion. This quantity is the product of *l0* and the residence time of the sample in laser beam Δ*t*, which is given by

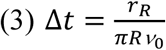

For a photochemically innocent probe beam, *l*_0_ · Δ*t* should be not larger than 0.1, such that the average fraction of the unphotolyzed state that is probed is ca. 95%. But even for a small photoconversion parameter, continuous rotation of the cell may lead to the accumulation of photocycle intermediates in the Raman probe beam if the rotational period of the cell (1/ν0) is too slow for a complete recovery of the photoconverted fraction of the sample. As shown in the Supporting Information (Fig. S2, Table S1), the ratio of the rotational frequency of the cell ν0 and the rate constant of the rate-limiting step of the photocycle *k*RL, which in BR is *k*O (Fig. 1A), is the critical quantity. Only for *k_RL_*/*v*_0_ ≥ 4, the accumulation of intermediates during continuous cell rotation through the laser beam is negligibly small, regardless of the extent of photoconversion during the residence time Δ*t* of the sample in the laser beam (Supporting Information, Fig. S2).

Selection of appropriate experimental parameters, therefore, should start by choosing the upper limit of the rotational frequency which, according to the *k_RL_*/*v_0_* ≥ 4 criterion, is 50, 25, 10, and 5 s^−1^ for photocycle recovery times of 5, 10, 25, and 125 ms, respectively. Then Δ*t* should be kept as small as possible, corresponding to the highest possible value for *ν*_0_ (Eq. 3), and *l*_0_should be adjusted as a tradeoff between low intermediate accumulation in the probe beam and an acceptable signal-to-noise ratio of the RR spectrum. The data in the Supporting Information (Table S1 and Fig. S1) may guide the choice of the experimental parameters. It can be seen that for *k_RL_*/*v*_0_ = 200 *s*^−1^ in confocal measurements *ν*_0_ should be between 25 and 50 s^−1^ and the laser power should not exceed 0.1 mW. For measurements with slit spectrometers, the limits for *ν*_0_ are the same but the acceptable maximum laser power is ca. 3 mW.

Whereas the two key criteria for probe-only RR spectroscopy can readily be fulfilled for BR, retinal proteins with photocycle periods longer than 100 ms require additional measures to achieve fresh sample conditions. In one technique, mechanical choppers may be used to increase the dark time.^[30]^ As a more versatile option, previously applied for sensory rhodopsin II, the cw laser beams were gated by opto-electronic shutters (Fig. 3).^[53]^ This approach allows blocking the laser beam for a desired period of time (i.e. *n* rotations of the cell) that is sufficient for the photoreceptor recovery (*n* · *k_RL_*/*v*_0_ ≥ 4). It is also applicable in two-beam pump-probe experiments. The disadvantage of this technique is a significant increase of spectra accumulation time. As an alternative one may employ the mixing-ball technique (Fig. 3).^[37,53]^ Here a magnetic ball is inserted into the cell and fixed at the inner wall of the rotating cell via a magnet, downstream of probe (and pump) lasers. This causes mixing of the irradiated volume element with the remainder of the solution before one rotation is completed. This corresponds to an effective increase of dark time *1*/*v*_0_ theoretically by factor of *V_cell_*/*V*_irr_ in case of perfect mixing, where *V*cell is the solution volume of the cell (here ca. ≤ 0.94 mL) and *V*irr the irradiated volume element (here ca. 1.0·10^−10^ mL). The real increase of the dark time is certainly smaller but still sufficient for probing photoreceptors with recovery times of many seconds. The disadvantage of this approach is the strong increase in shearing force which may cause degradation of the protein.

**Figure 3.**
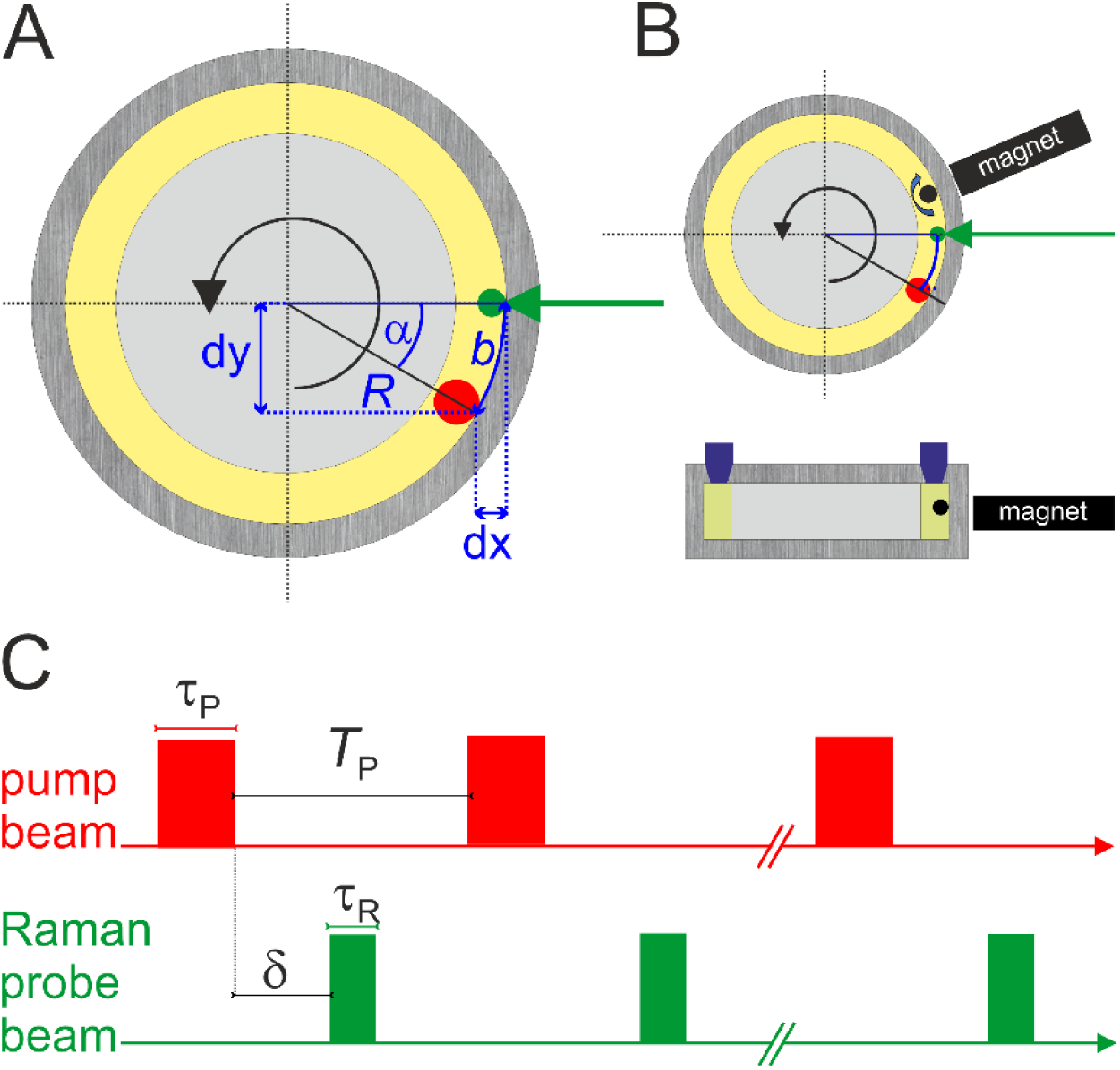
**A**, **B**. Schematic representation of the principles of pump-probe experiments using the rotating cell. **A**, top view, shows the displacement of the cw-probe (green) and -pump (red) beam with respect to each other. **B** illustrates the magnetic ball technique in a top view (top) and side view (bottom). **C** is a schematic representation of the gated-cw technique showing the gating of the pump and probe beams along the time axis.^[53]^ *T*p is the dark time between the pump events and δ denotes the delay time between pump and probe event. The duration of the pump and probe pulses are given by τ_P_ and τ_R_, respectively.

### 3.2. Pump – probe experiments

In pump-probe experiments the delay between photolysis and measurement event is defined by the beam separation Δ*s* and the velocity of the sample flow *v* (Fig. 1). For a rotating cuvette, the circular movement must be considered and Δ*s* is equal to the arc defined by the centers of the pump and probe beam passing through the cell (Fig. 3). The arc is given by

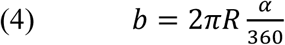

and related to the delay time δ by

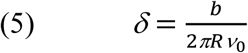

Taking into account (see Fig. 3A) that

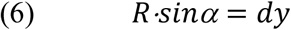

one may adjust the desired delay time by moving the pump beam in y- and x-direction according to

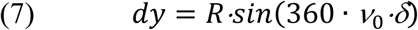

and

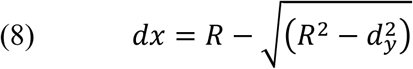

These relationships hold for both setups (Fig. 2). Whereas further details about pump-probe experiments with 90-degree-scattering geometry were described in detail previously,^[7,28,29,31,53,59]^ we now focus the particular challenges of the measurements with confocal spectrometers in backscattering (180 degree) geometry.

In general, the overall optical layout is optimized to keep the number of degrees of freedom for the alignment as small as possible, both for measurements and determining the focus size of the of pump beam (Fig. 4). Nevertheless, proper alignment of the pump and probe laser beams and adjustment of the experimental parameters is challenging in the confocal setup. This is due to the fact that the geometric shapes and orientations in the coordinate system of the volume elements irradiated by the probe and pump beams are quite different (Figs. 2, 4, 5). In the confocal setup, the pump laser beam passes through the cell in z-direction at 90 degree with respect to the Raman probe beam (x-direction). Its focus must be relatively large to facilitate alignment and to ensure that the photolyzed sample element runs through the focus of the Raman probe beam (Fig. 4, 5).

**Figure 4.**
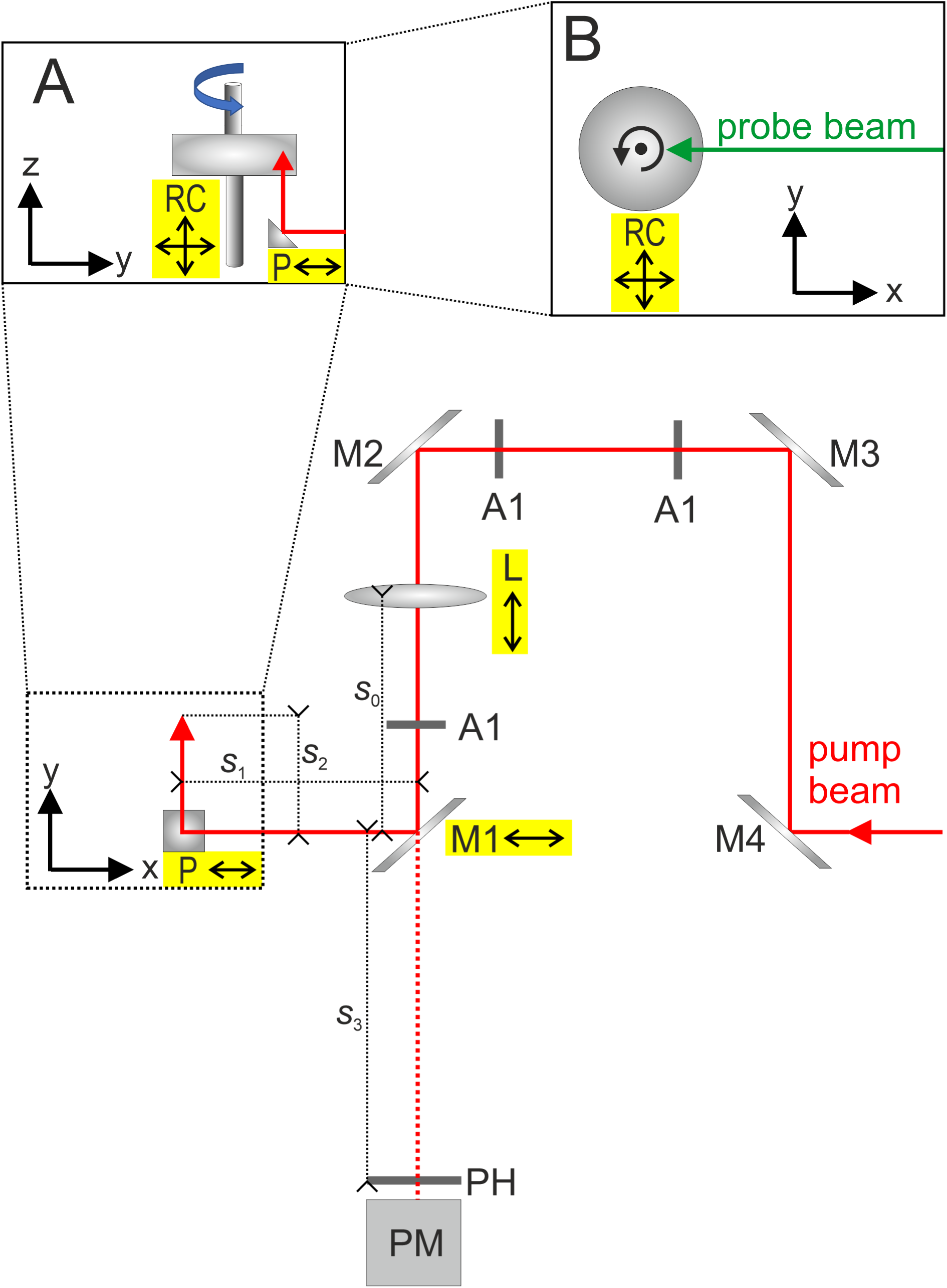
Optical layout of the pump-probe experiment with a confocal setup. Mirrors, apertures, lenses, and prisms are denoted by M, A, L, and P, respectively. Further abbreviations are PH, PM, and RC for the pinhole, power meter, and rotating cell, respectively. The arrows at the optical elements indicate the directions of alignment within the coordinate system as indicated.

**Figure 5.**
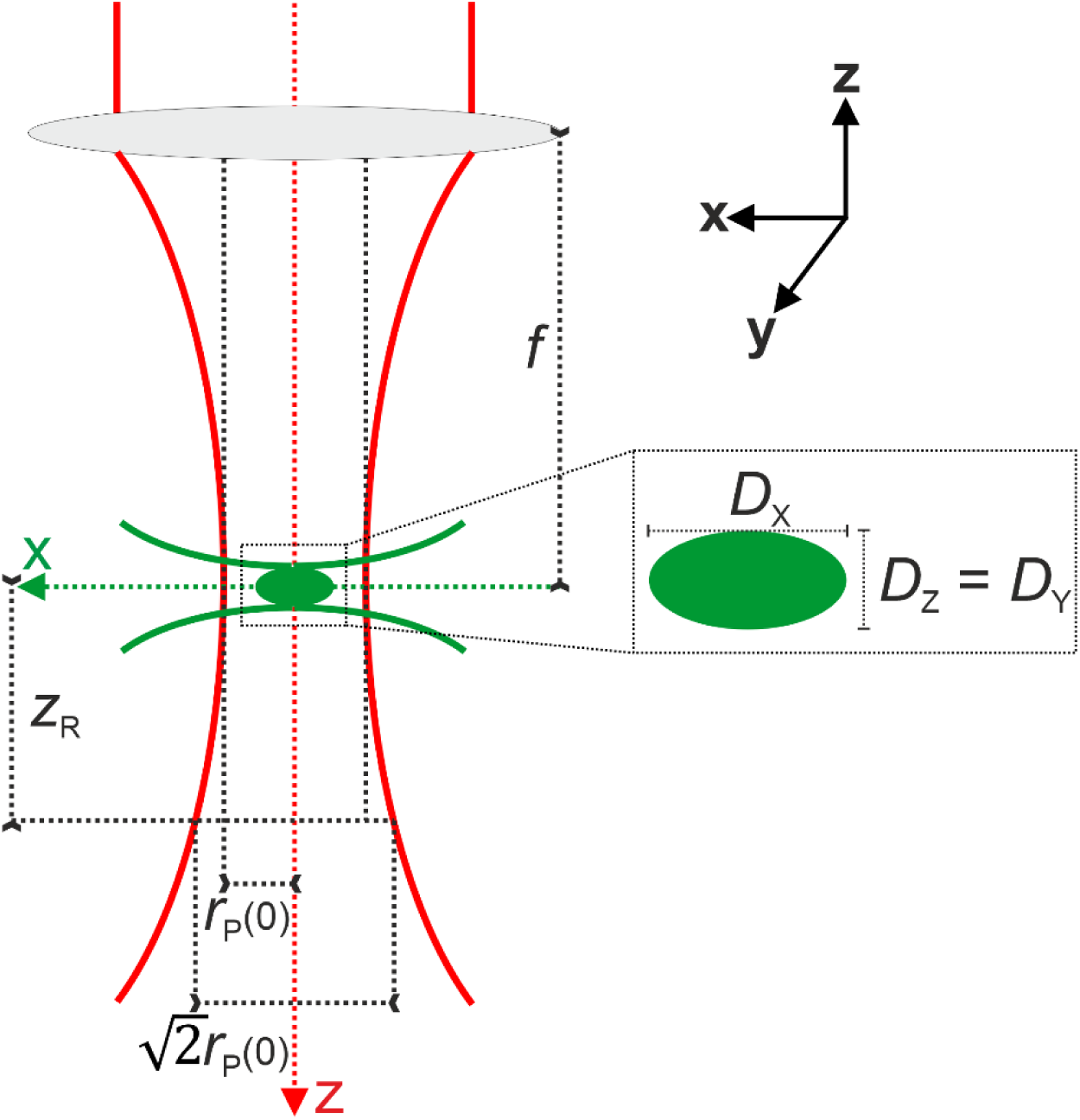
Sketch of the beam profiles of the pump and probe laser beams in the confocal set-up (see Fig. 4). The pump beam radius in the focus is denoted by *r*_P_(0) and the Rayleigh length is abbreviated by *z*_R_. The green ellipsoid shape shows the focus of the probe beam.

For the measurements, the pump beam is reflected by mirror M1 and the prism P to pass through rotating cell (RC), with the position of L adjusted to superimpose the focus of the pump beam in z-direction with the focus of the probe beam. The focal length *f* of L thus corresponds to *f* = *s*_0_ + *s*_1_ + *s*_2_. To determine the pump beam focus, mirror M1 is moved in *x*-direction out of the laser pathway such that the beam follows the dotted route through the pinhole PH. Here *s*3 is adjusted to *f* = *s*_0_ + *s*_3_such that the focus of L is in the pinhole. We can then determine the pump beam focus size at the sample as follows. The laser power *P* is related to the intensity of the laser light *I* as a function of the beam radius *r* and the propagation direction *z* according to

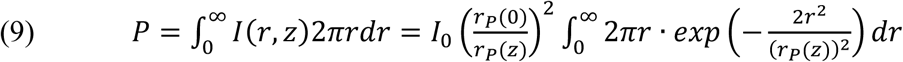

where *I*_0_ is the intensity at the beam waist center, *r*_*P*_(0) the waist radius in the focus, and *r*_*P*_(*Z*) the beam radius at the position where the intensity has dropped to *I*_0_/*e*^2^(Fig. 5). Upon solving the integral, Eq. 9 can be then rewritten to

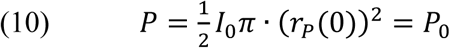

where *P*_0_ is the total laser power measured by the power meter. If we now consider the laser power *P*_*t*_ measured after transmission through the pinhole, we obtain – in analogy to Eq. 9,

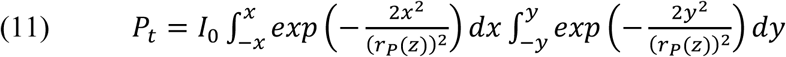

and with Eq. 10 and for *x* = *y* and *r*_*P*_(*Z*) = *r*_*P*_(0)

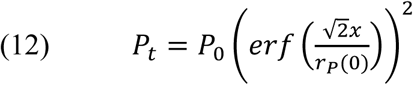

The focus diameter 2*r*_*P*_(0) can thus readily be obtained by measuring *P*_*t*_ via a pinhole with diameter 2·*x* (Eq. 12) and *P*_0_, and using tabulated values of the error function.

The beam radius as a function of the distance *d* from the lens L is evaluated by

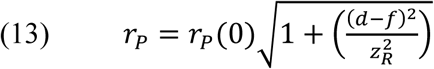

where *z*R is the Rayleigh length given by

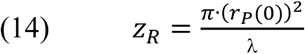

The precise positioning of the pump beam with respect to the probe beam is critical for the success of the experiments. The focus of the probe beam forms a spheroid that can be approximated by an Airy disk.^[60,61]^ In *z*- and *y*-directions the diameters are the same and given by

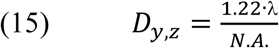

where *N.A.* is the numerical aperture. For the diameter in *x*-direction, Eq. 16 holds

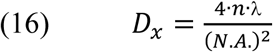

with *n* denoting the refractive index of the sample. The sample volume *V*_*R*_ that is irradiated by the probe beam is then approximated by

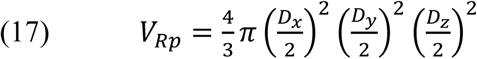

For a numerical aperture *N.A.* = 0.35, a refractive index of *n* = 1.5, and a laser wavelength of 514.5 nm, the values for *D*_*y,Z*_ and *D*_*x*_ are 1.79 and 25.2 μm, respectively. The different shapes of the irradiated volume elements of pump and probe beam impose some difficulties for the theoretical description and the alignment of the beams, as compared to the setup for slit-spectrometer. In the latter case, the propagation directions of pump and probe beam are identical and also the shapes of the irradiated volume elements qualitatively the same and sufficiently characterized by the focal planes π · *r*^2^ and π · *r*^2^. For the theoretical treatment in the confocal setup, i.e. the evaluation of *l*_0_ as a function of the laser power, we therefore must translate the irradiated volume of the Raman probe beam into an effective irradiated area with a radius *r*R derived from a sphere with a volume of *V*_*R*_ (Eq. 17). Accordingly, we obtain a value of ca. 2 μm for excitation lines in the visible range.

As a consequence, the focal waist of the pump beam must be distinctly larger than 25 μm to allow for a reliable superposition of the pump- and probe-irradiated volume elements. Large focal diameters of the pump beam of more than 100 μm, however, reduce the photon flux and thus the photoconversion parameter *l*_0_ · Δ*t*, which for the pump beam should be as large as possible. The photoconversion parameter is further reduced due to the strong absorption of pump laser intensity upon passing through the cell in z-direction. Unlike for 90-degree-scattering geometries with optimum concentrations corresponding to ca. 1 OD, backscattering (180-degree scattering) geometry requires distinctly higher concentrations (ca. ≥ 5 OD) to achieve a good signal-to-noise (S/N) ratio, leading to a sharp gradient of photoconversion in z-direction. As a consequence, the common volume element that is photolyzed and probed should be as close as possible to the wall where the beam enters the cell. Such an alignment bears the risk of reflections from the cell wall which produces strong stray light and thus lowers the S/N ratio.

### 3.3. Choosing experimental parameters for time-resolved RR experiments

As shown in the previous sections, proper TR RR measurements of retinal proteins or photolabile molecules in general is not trivial, and requires careful selection of the experimental parameters. This section is dedicated to support the planning of the experiment by pointing out the most critical issues.

In the simplest case, the target state is the measurement of the parent dark state in a probe-only TR RR experiment. Here, the main criteria, the photochemical innocence of the probe beam and the fresh sample condition, have to be fulfilled. The fresh sample condition is fulfilled for *k_RL_*/*v*_0_ ≥ 4, which for a given probe beam and cell radius is solely given by the rotational frequency of the cell (see Eq. 3; Supporting Information, Fig. S2). For BR and the related retinal protein HR, which exhibits similar kinetic and spectral properties, the rate-limiting steps of the photocycle *k*_*RL*_ are between 100 and 200 s^−1^.^[2,55,62]^ Thus, the fresh sample criterion can readily be satisfied using rotational frequencies between 20 and 50 s^−1^. If for other systems the recovery time of the sample is too low or in the limiting case of irreversible processes, additional measures like the magnetic-ball technique or the gated cw excitation have to be employed (see Fig. 1). The wavelength of the probe beam must be selected as a compromise between optimum resonance conditions and low photochemical activity. To reduce photoconversion of the sample in the probe beam, the laser power should be as low as possible while still maintaining an acceptable spectra acquisition time (see Eq. 2; Supporting Information, Fig. S2). In principle, also intermediate spectra can be obtained in single beam experiments by comparing spectra of low and high laser power and/or short and long residence time of the sample in the laser beam. This approach, however, lacks the possibility of choosing the kind of intermediates by adjusting the delay time and, in addition, bears the risk of generating secondary photoreactions.

An accurate TR RR procedure to measure intermediate states and determine the reaction kinetics of the system requires two-beam experiments. The choice of the pump beam, its positioning and properties, depends on the settings for the Raman probe beam as discussed above. For probing different states, it is desirable that the probe wavelength provides comparable resonance enhancement for the parent state and all intermediates under consideration. Nevertheless, due to the different RR cross sections, one cannot directly equate relative spectral contributions with relative concentrations of the various species. Instead, approximations may be employed as discussed below, or as a more accurate method, one may use an internal standard such as sulfate or perchlorate. However, in case of proteins, which are sensitive to the ionic strength, high salt concentrations may affect the kinetics.

The wavelength and the power of the pump beam must be selected to ensure efficient photoconversion. Wavelengths of the pump and probe beams should be spectrally separated to avoid interferences of the RR spectrum by the stray light and RR scattering due to the pump beam (Supporting Information, Fig. S3). For the experiments with BR and HR, we have used 488 and 514 nm probe wavelengths and 568 and 647 nm pump wavelengths.

Finally, the appropriate sample concentration must be chosen as a tradeoff between good S/N ratios and low photoconversion on the one hand, and long accumulation times and good photoconversion on the other hand. Thus, we have used a concentration of ca. 150 μM for the confocal set-up whereas in the slit-spectrometer experiments the concentration was between 15 and 25 μM.

### 3.4. Time-resolved spectroscopy of BR using a confocal spectrometer

In view of the recovery time of 7 ms, we chose a rotational frequency of 20 s^−1^. The probe wavelength was 488 nm with a laser power of 0.2 mW such that the average contribution of the parent state in the probe beam was ca. 95%. Typical total accumulation time, corresponding to the acquisition time multiplied by the number of repetitions, was ca. 90 min. The pump wavelength was 568 nm with a power of ca. 100 mW at the sample.

The probe-only spectrum of the parent state BR570 is of high quality (Fig. 6). It agrees very well with previously published spectra, including those recorded with slit-spectrometers as shown below.^[6,7]^ A careful inspection of the originally measured spectrum revealed a very weak band of the L550 intermediate that is formed during the probe event. With respect to the strongest bands (C=C stretching modes), the spectral contribution was less than 3% confirming the above estimate from the photoconversion parameter. In Fig. 6, we have subtracted the small contribution from the measured probe-only spectrum.

**Figure 6.**
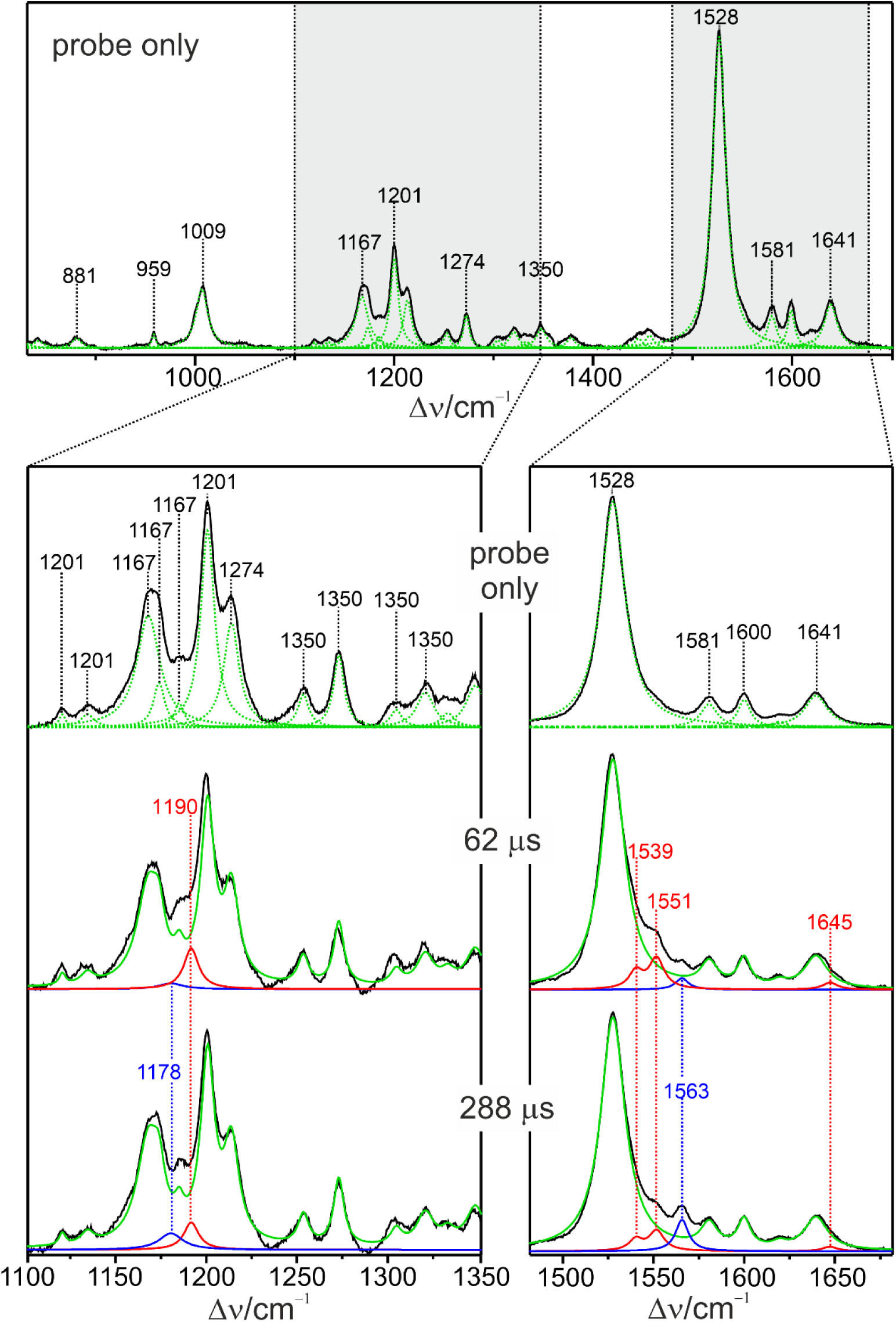
TR RR spectra of BR measured with a confocal spectrometer and 488 nm excitation. The probe-only spectrum is compared with the pump-probe spectra obtained with delay times of 62 and 288 μs, reflecting rise and decay of the L550 intermediate. The spectra were subjected to a component analysis with the spectra of BR570, L550, and M410 shown in green, red, and blue, respectively. Spectral regions that a diagnostic of the retinal configuration and Schiff base protonation state are highlighted in gray (top) and shown in a close-up view at the bottom. The total accumulation time for each spectrum was 90 min.

Pump-probe experiments were carried out with variable delay times from 12 to 1276 μs, such that contributions from the intermediates L550 and M410 are expected. The spectral changes were relatively small as shown by two representative pump-probe spectra. Specifically, a weak shoulder at 1551 cm^−1^ and, at longer delay times, an additional band at 1563 cm^−1^ were observed.

These bands could readily be attributed to the intermediates L550 and M410. A component analysis of the spectra,^[52]^ using the BR570 spectrum (probe-only) as a fixed fit component, indeed identified two intermediate spectra with the prominent bands at 1190, 1539, 1551, and 1645 cm^−1^ as well as 1178 and 1563 cm^−1^, constituting the component spectra of L550 and M410, respectively. All experimental pump-probe spectra could then readily be simulated by a superposition of the three component spectra. The amplitudes of the component spectra of L550 (red solid circles) and M410 (blue solid circles) were then plotted as a function of the delay time (Fig. 7).

**Figure 7.**
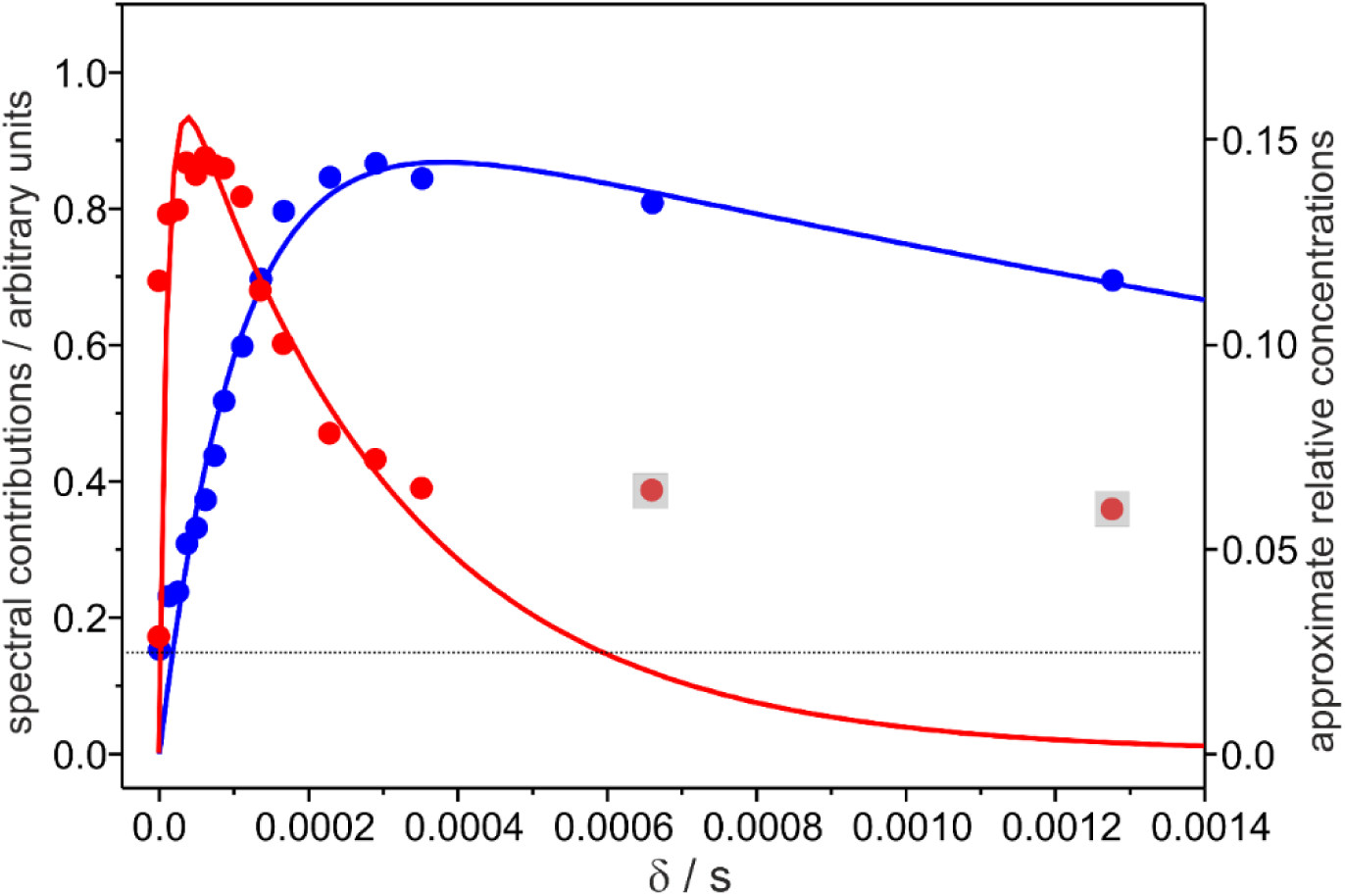
Relative contributions of the component spectra (spectral contributions; scale on the left ordinate) of the L550 (red) and M410 (blue) intermediates as a function of the delay time in pump-probe experiments using the confocal spectrometer. The solid lines represent exponential fits to the data points. Here, the gray-shaded data points were not considered for the fit. The horizontal dotted line marks the upper limit of L550 contributions produced by the probe beam. The scale on the right ordinate refers to the approximate relative concentrations assuming that at 488 nm the resonance enhancement of the C=C stretching modes are the same for the intermediates and BR570.

To convert the amplitudes of the component spectra to the relative concentrations requires the relative RR cross sections of the three species at 488 nm excitation. These quantities are not known a priori but, as a first approximation, one may assume that at 488 nm the resonance enhancement of the C=C stretching modes are the same for the intermediates and BR570. This approximation, which does not affect the rate constant determination, presumably over- and underestimates the L550 and M410 species, respectively. Thus, the maximum extent of photoconversion of ca. 15% of the total protein is certainly only a crude estimation. Nevertheless, it allows for comparison with the corresponding values from the spectra measured with the slit-spectrometer set-ups, where the same approximation was applied (vide infra).

The temporal evolution of L550 and M410 was separately described by exponential functions yielding rate constants for the formation and decay of L550 and M410, with an acceptable average error of ca. 17% (Table 1).

**Table 1.** Rate constants of the photocycle of BR derived from TR RR spectroscopy^a^.

| rate constant | this work |  | literature<sup>b</sup> |
| --- | --- | --- | --- |
|  | L550 data<sup>a</sup> | M410 data<sup>a</sup> |  |
| K610 → L550, $k_K$ | $9.1 \cdot 10^4 \text{ s}^{-1}$ | | $1 \cdot 10^6 - 4 \cdot 10^6 \text{ s}^{-1}$ |
| L550 → M410, $k_L$ | $3.4 \cdot 10^3 \text{ s}^{-1}$ | $9.4 \cdot 10^3 \text{ s}^{-1}$ | $2 \cdot 10^4 - 1 \cdot 10^5 \text{ s}^{-1}$ |
| M410 → N560, $k_M$ | | $2.9 \cdot 10^2 \text{ s}^{-1}$ | $2 \cdot 10^2 - 3.5 \cdot 10^2 \text{ s}^{-1}$ |

<sup>a</sup> The rate constants were determined separately for the individual data sets, based on the simplified reaction scheme in Fig. 1. The average error was $\pm 17\%$ .
<sup>b</sup> data reflect the range of values reported in the literature for pH 7, T = 20 °C (see cited studies and references therein).<sup>[3,63,64]</sup> The rate constants are model-dependent.

However, there are severe discrepancies particularly related to the data for L550. At long delay times, the relative concentrations of L550 do not decay to zero but seem to approach a constant value of ca. 5%. This can only partly be attributed to the formation of L550 in the probe beam (ca. 2.5%). A more critical factor is the simple consecutive reaction scheme that constitutes the basis for the present analysis. This scheme ignores multiple L550 (sub-)states, possible secondary photoreactions of L550 in the probe beam, formation of N560, reaction equilibria, as well as parallel reaction cycles.^[3,25,28,31,32,63–68]^ Employing a more complex reaction scheme for the analysis of the present spectra, however, is not justified in view of the very small relative concentrations of the intermediates in the pump-probe experiments that do not exceed ca. 15%. Under these conditions, spectrally slightly different L550 substates (i.e., early and late L550)^[28]^ cannot be identified, particularly since the characteristic vibrational bands of L550 strongly overlap with those of the dominant parent state BR570. Both the low extent of L550 formation and the simplified reaction scheme may therefore also account for the discrepancy of the *k*L values derived from the L550 and M410 data, as well as the strong deviations of *k*K and *k*L from literature data (Table 1). Conversely, the value for the M410 decay *k*M falls into the range of literature values, evidently since, despite a structural heterogeneity also for the M410 state, the overall decay to N560 can be approximated by a single exponential function.

### 3.5. Time-resolved spectroscopy of BR and HR using a slit spectrometer

Important contributions to the elucidation of the photocycle mechanism and kinetics of BR were provided by the TR RR measurements of the Stockburger group.^[28,31,32]^ The experiments were carried out with a slit-spectrometer set-up using a scanning double monochromator, equipped with a photomultiplier and a two-channel photon-counting device. The latter allowed for quasi-simultaneous measurements of probe-only and pump-probe events. This was achieved by repetitively blocking the pump beam and feeding the signals of pump-on and pump-off into separate photon counting channels. Thus, regardless of the very long data acquisition time of many hours, a high frequency precision of the conjugate pump-only and pump-probe spectra was guaranteed, as a prerequisite for an accurate quantitative analysis of the spectra (Fig. 8). The second important factor was the relatively high extent of photoconversion by the pump beam which led to relative concentrations of L550 of more than 30%. In contrast, the L550 content produced by the probe beam was kept sufficiently low with less than 5%.

**Figure 8.**
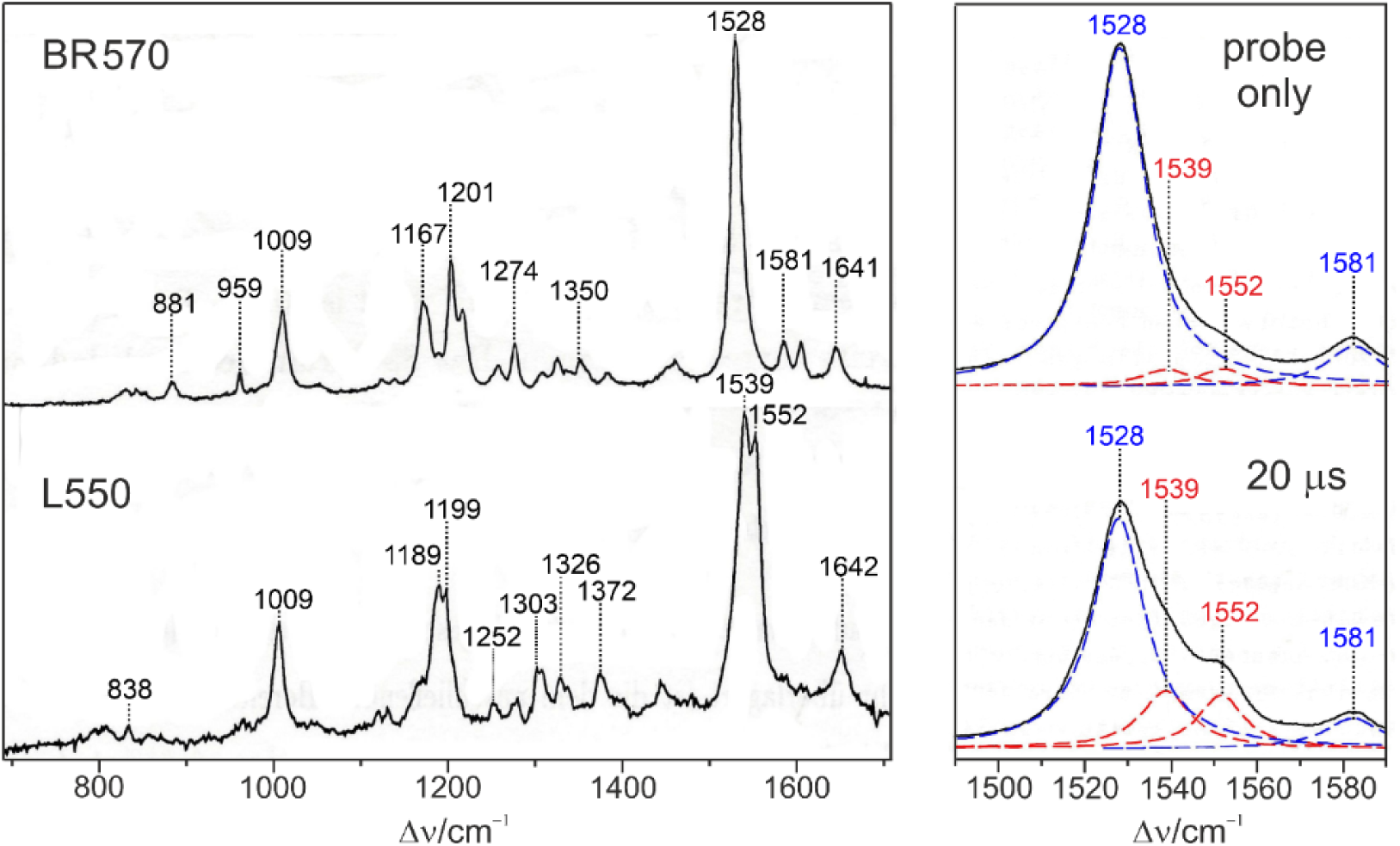
TR RR spectra of BR measured with a scanning double monochromator and 488 nm excitation.^[28,69]^ The left panel shows the spectra of BR570 (top) obtained in a probe-only experiment, and L550 (bottom), measured in a pump-probe experiment (pump wavelength 647 nm, 600 mW) with a delay time of 20 μs, followed by subtraction of the residual BR570 content. Subtraction was guided by the band fitting of the C=C stretching region that is shown in a close-up view in the right panel with the bands of BR570 and L550 in blue and red, respectively. Note that the spectra in the right panel refer to the same intensity scale reflecting the increase of the L550 intermediate at the expense of the parent state (bottom) compared to the probe-only spectrum (top). The total accumulation for each spectrum was ca. 18 hours.

The major disadvantage of scanning detection systems is the long accumulation time. This drawback can be improved using a slit spectrometer operated as a spectrograph and equipped with a CCD detection system. Thus, the advantage of the 90-degree scattering geometry, i.e. the high extent of photoconversion, is preserved whereas the accumulation time required for an acceptable S/N ratio is significantly reduced. As an example, Figure 9 shows the probe-only and pump-probe spectra of HR of *Natronomonas pharaonis* in detergent, which exhibits very similar kinetic and spectral properties as BR.

**Figure 9.**
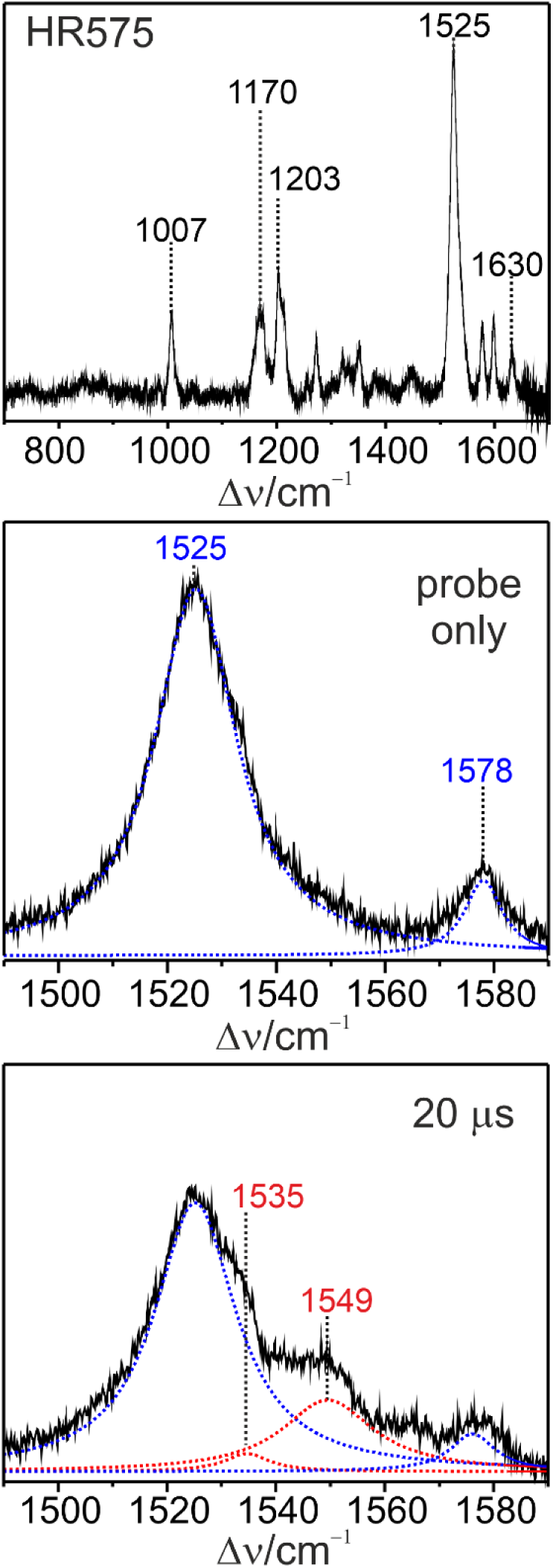
TR RR spectra of HR from *Natronomonas pharaonis* measured with a double monochromator spectrometer, operated as a spectrograph with CCD detection, and 488 nm excitation. The probe-only spectrum of HR575 is compared with the pump-probe spectrum (pump wavelength 647 nm, 600 mW) obtained with a delay time of 20 μs, reflecting the formation of the HR520(L1) intermediate (Supporting Information, Fig. S3). The spectra were analyzed by a band fitting program, with the bands of HR575 and HR520(L1) shown in blue and red, respectively. Note that the spectra refer to the same intensity scale reflecting the increase of the HR520(L1) intermediate at the expense of the parent state (bottom) compared to the probe-only spectrum (top). The top spectrum included less than 2% of HR520(L1), not considered by the band fitting. The total accumulation time was ca. 10 min and thus by factor of nearly 10 and 100 shorter than for the spectra in Fig. 6 and 8, respectively.

### 3.6. Comparison of the different set-ups for time-resolved resonance Raman spectroscopy

We now compare the performance of the three set-ups on the basis of selected parameters that are relevant for TR RR spectroscopy of retinal proteins (Table 2).

**Table 2.** Performance of three set-ups for time-resolved RR spectroscopy.

| Parameter<sup>a</sup> | confocal spectrometer, CCD<sup>b</sup> | slit spectrograph, CCD<sup>c</sup> | slit spectrometer, photon counting<sup>d</sup> |
| --- | --- | --- | --- |
| spectral resolution | 2 cm<sup>-1</sup> | 2 cm<sup>-1</sup> | 4 cm<sup>-1</sup> |
| wavenumber increment per pixel or step width | 0.15 cm<sup>-1</sup> | 0.15 cm<sup>-1</sup> | 0.2 cm<sup>-1</sup> |
| $T_{ac,50}$, acquisition time for S/N = 50 | 10 min | 30 min | 350 min |
| protein, sample concentration | BR, 150 μM | HR, 20 μM | BR, 15 μM |
| photoconversion (max. intermediate concentration) | 15% | 30 - 35% | 30 - 35% |

<sup>a</sup> The values for the individual parameters correspond to the experimental conditions of the spectroscopic measurements reported in this work.
<sup>b</sup> Horiba HR-800 confocal spectrometer
<sup>c</sup> ISA, U1000 spectrograph
<sup>d</sup> Spex 14018 double monochromator,

For a quantitative analysis, a spectral resolution of 4 cm^−1^ of the TR spectra is sufficient which can readily be achieved in all three set-ups. This is also true for the wavenumber increment per CCD pixel, corresponding to the step width in the scanning detection, with desirable values of 0.2 cm^−1^ or less. The main difference between the three set-ups is related to the data acquisition time *T*ac required to achieve a high S/N ratio. For the comparison of the set-ups, we take a S/N ratio of 50 with respect to the strongest RR band as a criterion for a very good spectral quality. The acquisition times *T*ac,50 refer to the spectra shown in this work and measured with sample concentrations indicated in Table 2. The actual values of *T*ac were extrapolated to S/N = 50, taking into account that S/N scales with the square root of *T*ac. The quantities *T*ac,50 depend on various factors such as optical alignment and light collection or quality of the sample, which may vary from lab to lab and experiment to experiment. Therefore, the *T*ac,50 values listed in Table 2 are crude estimates to illustrate the qualitative differences between the three set-ups.

For the 90-degree scattering geometry, the ideal sample concentration corresponds to an optical density of 1 for the pathway (1 cm) of the probe beam through the rotating cell. This concentration is also ideal for an efficient photoconversion. The value of 30 - 35% photoconversion is close to the maximum possible value in view of a photochemical quantum yield of ca. 0.67,^[57]^ and the photochemical back reaction in the pump beam. In the confocal set-up, the concentration should be much higher in probe-only experiments in order to achieve high signal intensities, which on the other hand reduces the extent of photoconversion in pump-probe experiments. Thus, sample concentration represents a trade-off between signal quality and product formation in confocal set-ups.

High relative concentrations of the intermediates are necessary for a reliable quantitative analysis of the TR RR spectra. The analysis further requires removal of the large spectral contribution of the unphotolyzed reference state from the pump-probe spectra, either by direct spectra subtraction or via component analysis. In any case, identical frequency calibration of probe-only and pump-probe spectra is essential, which is a challenge for long-term measurements. Here, the scanning system with a dual-channel photon counting system offers an ideal solution which allows for quasi-simultaneous measurements of probe-only and pump-probe spectra. However, this advantage is essentially paid by doubling the already rather long accumulation times.

## 4. Conclusions

In this work, we have recalled essential principles of TR RR spectroscopy of retinal proteins. These principles were developed decades ago for spectrometer and detection systems that have meanwhile been largely replaced by more efficient devices, usually confocal spectrometers. It was shown that the slit spectrometers had certain advantages for this specific application. However, we are aware that nowadays researchers in this field only rarely have the opportunity to choose between different set-ups. Therefore, the main objective was to show a possible approach to adapt the critical principles of TR RR spectroscopy to confocal spectrometer set-ups. In the present study, this adaptation was accomplished using a rotating cell, which appears to be the most convenient sample device. Unfortunately, rotating cells as used in this work are nowadays not easily available but must be manufactured upon request. NMR or EPR tubes, frequently used as alternatives, have a distinctly smaller radius such that the residence time in the laser beam increases unless the rotational frequency is raised correspondingly. An increase of ν0, however, may be in conflict with the “fresh sample” condition. Thus, the TR experiments with such tubes would most likely require the use of gated cw pumping and probing.^[53]^ Another option is a capillary flow system which, however, limits high protein concentration due to the increased viscosity, and shearing forces may lead to protein foaming and denaturation.

## Acknowledgements

We thank Peter Hegemann and his team for providing the BR samples. The work was supported by the Einstein Stiftung Berlin EVF 2018-427 (to K.F. and P.H.) and the Deutsche Forschungsgemeinschaft (DFG) through grants SFB1078 “Protonation Dynamics in Protein Function”, project number 221545957, subproject B6 (to P.H.), and the DFG under Germany’s Excellence Strategy—EXC 311 2008/1 (UniSysCat)—390540038, Research Unit E (to P.H.).

## Data Availability Statement

The data that support the findings of this study are available from the corresponding author.

## Dedication

We dedicate this study to the memory of Manfred Stockburger and to Richard A. Mathies on the occasion of his 80th birthday (2026).

## Graphical Table of Content

We have adapted pump-probe time-resolved resonance Raman spectroscopic experiments with continuous wave excitation to a confocal spectrometer. Based on the photocycle of bacteriorhodopsin and the rotating cell technique, we analyse the parameters to maintain fresh-sample condition and photochemical innocence of the probe beam while increasing the extent of photoconversion in the pump beam. The performance of the experiments is critically compared with the results obtained with classical slit spectrometers.

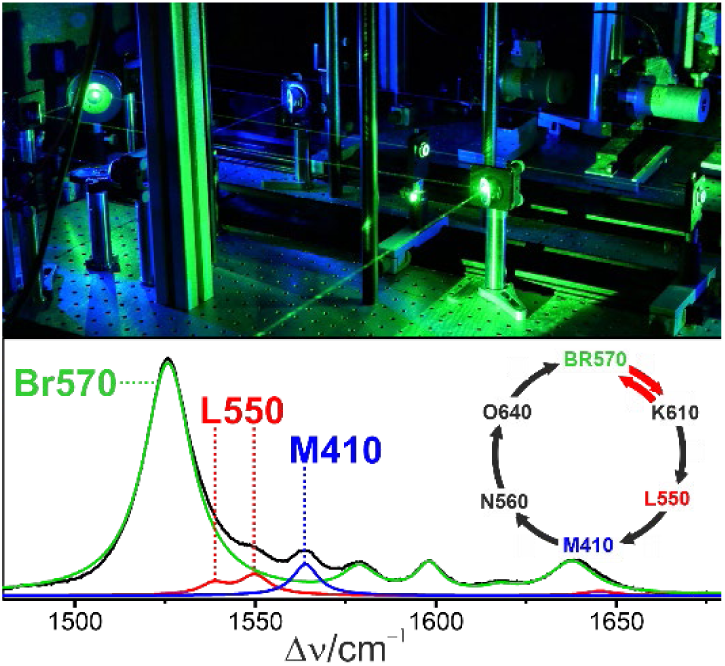

