## Supporting Information for "Time-Resolved Resonance Raman Spectroscopy of Retinal Proteins with Continuous-Wave Excitation. A Fundamental Methodology Revisited"

### 1. Photoconversion in the Raman probe beam and accumulation of intermediates

In the following, we consider a model photoreceptor, denoted as BR for the sake of simplicity although specific spectral and kinetic properties are not exactly the same as in bacteriorhodopsin. The extent of photoconversion of the sample flowing through the laser beam is given by the residence time of the sample in the beam Δt and the photochemical rate constant $l_0$. Here we restrict the discussion to a rotating cell with a radius of R = 13 mm with a rotational frequency $\nu_0$ that may range between 5 and 50 $s^{-1}$. Then the residence time is given by

(S1) $$\Delta t = \frac{r_R}{\pi R \nu_0}$$

where $r_R$ is the radius of the laser beam, here the Raman probe beam. Here and in the following, the laser beam radius is defined by the drop of the laser intensity from its maximum value $I_0$ to $I_0/e^2$.

The photochemical rate constant for the conversion of parent state to the primary photoproduct within an irradiated volume element passing the laser beam, $l_0(t)$, is given by[1–3]

(S2) $$l_0 = \frac{ln10}{\sqrt{2\pi} \cdot h \cdot c \cdot N_A} \cdot \gamma \cdot \varepsilon(\lambda) \cdot \lambda \cdot \frac{P}{r_R^2} \cdot exp\left(-8\frac{t^2}{\Delta t^2}\right)$$

To convert Eq. (S2) into a time-independent equation, the exponential term is integrated from –∞ to +∞, and by inserting the numerical values for *h*, *c*, and $N_A$ one obtains

(S3) $$l_0 = 0.481 \cdot \gamma \cdot \varepsilon(\lambda) \cdot \lambda \cdot \frac{P}{r_R^2}$$

where $\gamma$, $\lambda$, and $\varepsilon(\lambda)$ denote the photochemical quantum yield, the wavelength of the Raman probe beam (in nm), and the extinction coefficient of the retinal chromophore at $\lambda$ (in $L \cdot mol^{-1} \cdot cm^{-1}$), respectively. For a probe wavelength of 514.5 nm, and $\gamma = 0.5$ and $\varepsilon(514) = 30000\ L \cdot mol^{-1} \cdot cm^{-1}$) that approximately hold for BR and many other retinal proteins,[4] Eq. (3) is given by

(S4) $$l_0 = 3.712 \cdot 10^6 \cdot \frac{P}{r_R^2}$$

with $l_0$ in $s^{-1}$, $P$ in mW and $r_R$ in µm. The photoproduct K may either thermally decay to the L intermediate with an approximated rate constant of $k_K$, or it converts back to BR photochemically ($l_B$), as depicted in Fig. 1 of the manuscript. Due to the red-shifted absorption maximum of K the corresponding rate constant $l_B$ is estimated to be three times smaller than $l_0$. In a first approximation we assume that these three rate constants determine the concentration changes of the sample in the Raman beam according to

(S5) $$\frac{d[BR]}{dt} = -l_0[BR] + l_B[K]$$

(S6) $$\frac{d[K]}{dt} = +l_0[BR] - (l_B + k_K)[K]$$

(S7) $$\frac{d[L]}{dt} = +k_K[K]$$

The solution of this set of differential equations is given by

(S8) $$[BR] = \frac{[C]_0}{2\lambda_0}\{D_1 exp(-\lambda_1 t) + D_2 \exp(-\lambda_2 t)\}$$

(S9) $$[K] = \frac{l_0[C]_0}{\lambda_0}\{exp(-\lambda_1 t) - \exp(-\lambda_2 t)\}$$

with

(S10) $$\lambda_0 = \sqrt{l_0^2 + l_B^2 + k_K^2 - 4l_0 l_B}$$

(S11) $$\lambda_1 = \frac{l_0 + l_B + k_K - \lambda_0}{2}$$

(S12) $$\lambda_2 = \frac{l_0 + l_B + k_K + \lambda_0}{2}$$

(S13) $$D_1 = \lambda_0 - l_0 + l_B + k_K$$

(S14) $$D_2 = \lambda_0 + l_0 - l_B - k_K$$

We assume that the total concentration of the protein $[C]_0$ remains unchanged and set $[C]_0 = 1$, the concentration of L is then given by

(S15) $$[L] = 1 - [BR] - [K]$$

This kinetic model neglects the photoreaction of L,[4] and thus overestimates [L] in the laser beam. We can therefore take the values calculated for [L] as upper limits.

On the basis of this model and using $k_K = 10^6\ s^{-1}$ we have determined the time-dependent concentration profile of BR during passage of the sample through the laser beam as a function of the laser power. We have considered two cases representing measurements with a confocal spectrometer with $r_R = 2\ \mu m$ and a slit spectrometer with $r_R = 30\ \mu m$. A selection of the results is plotted in Figure S1; more data is listed in Table S1.

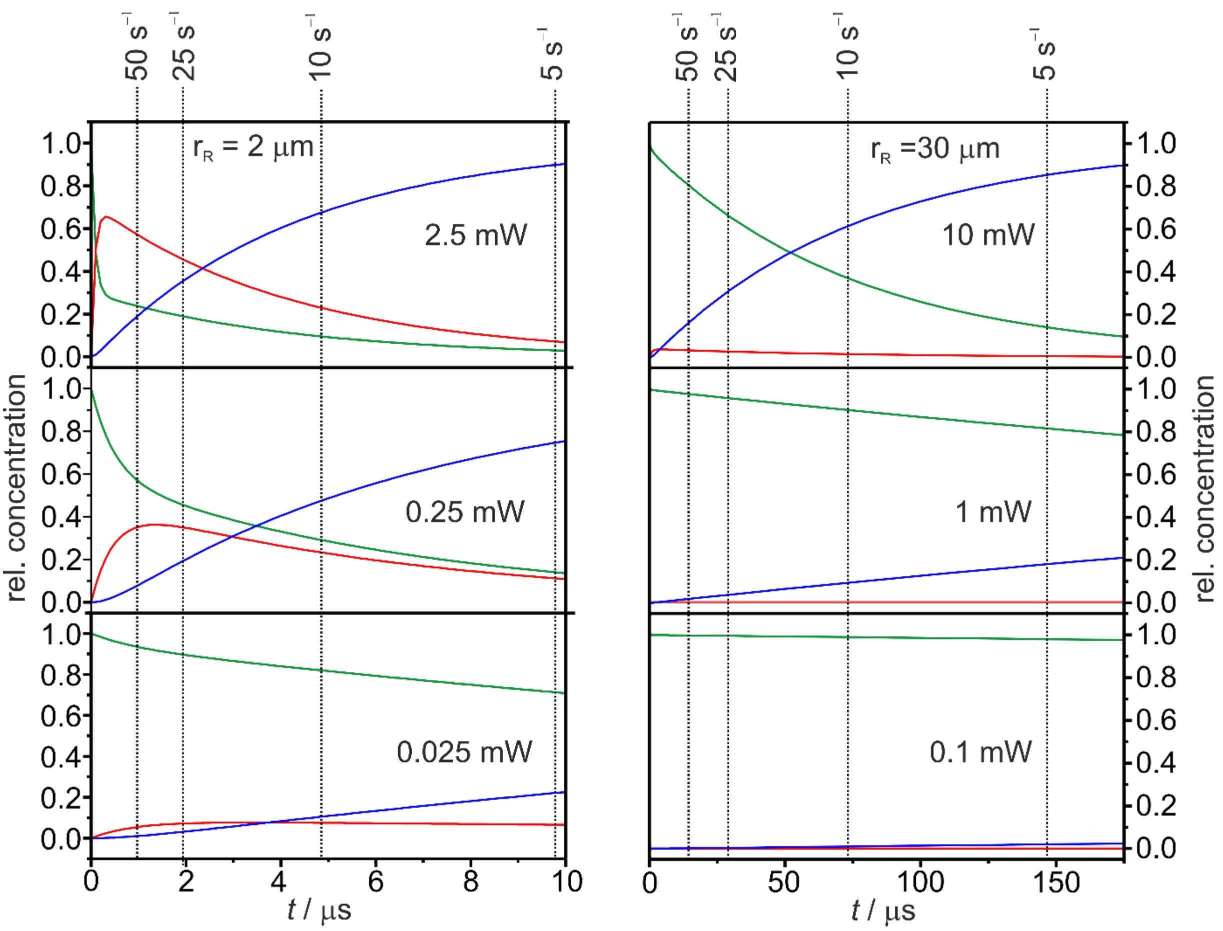


**Figure S1.** Temporal evolution of the concentrations BR (green), K (red), and L (blue) under the impact of $l_0$ (eq. S4) over time $t$ for different laser foci (left and right panel) and laser powers (top to bottom). The relative concentrations were evaluated according to the Eqs. S8 – S15 on the basis of the reaction scheme in Fig. 1 of the manuscript. The vertical dotted lines mark the residence times of the sample in the Raman probe beam for different rotational frequencies and laser foci.

The photoconversion parameter $\Delta t \cdot l_0$, i.e. the product of Eqs. S1 and S2, may be used to describe extent of photoconversion. As a good rule of thumb, $\Delta t \cdot l_0$ should be ≤ 0.1 for the photochemically innocent probe beam such that the fraction of the probed parent state BR is at average $0.5 \cdot ([BR]_{t=0} - [BR]_{t=\Delta t}) \geq 0.95$ during the probe event. In the present example of BR and 514.5 nm excitation, the photoconversion parameter is given by

(S16) $$\Delta t \cdot l_0 = 90.89 \frac{P_0}{r_R \nu_0}$$

For $r_R = 2\,\mu m$ (confocal spectrometer) appropriate conditions are $\nu_0 = 25\,s^{-1}$ and $P = 0.05\,mW$ whereas for $r_R = 30\,\mu m$ (slit spectrometer) $\nu_0 = 25\,s^{-1}$ and $P = 0.9\,mW$ represent ideal conditions.

To avoid accumulation of intermediates during the repetitive passage through the laser beam, the photocycle must be completed before an irradiated volume element of the sample re-enters the laser beam. If Δx denotes the fraction of the sample photoconverted after the passage through the laser, after one rotation of the cell the relative concentration of the parent state, Δ[BR], that reenters the laser beam, is given by $(1 - \Delta x)\exp\left(-\frac{k_{RL}}{\nu_0}\right)$, where $k_{RL}$ is the rate constant for the rate-limiting step of BR recovery, i.e. the slowest reaction of the photocycle. After the second rotation, Δ[BR] is then given by $1 - \left(\Delta x \cdot \exp\left(-\frac{k_{RL}}{\nu_0}\right) + \Delta x\right)\exp\left(-\frac{k_{RL}}{\nu_0}\right)$ and after the third rotation by $1 - \left(\left(\Delta x \cdot \exp\left(-\frac{k_{RL}}{\nu_0}\right) + \Delta x\right)\exp\left(-\frac{k_{RL}}{\nu_0}\right) + \Delta x\right)\exp\left(-\frac{k_{RL}}{\nu_0}\right)$ which is continued as a geometric series. The limiting value of Δ[BR] is then given by

(S17) $$\Delta[BR]_{lim} = 1 - \frac{\Delta x \cdot \exp\left(-\frac{k_{RL}}{\nu_0}\right)}{1 - \exp\left(-\frac{k_{RL}}{\nu_0}\right)}$$

The results are listed in Table S1 and shown in Figure S2. Note that the calculation described above does not consider a displacement of the irradiated sample element out of the laser focus due to diffusion or centrifugal forces. One may estimate that on the time scale of ass rotational period of the cell such a displacement is ca. $10^{-3}$ to $10^{-4}$ % of the laser focus radius and thus can be neglected.[5] In Figure S2, $\Delta[BR]_{lim}$ is plotted as a function of the logarithm of $l_0$ for various $k_{RL}/v_0$ ratios. It is evident that regardless of the degree of photoconversion in the laser beam, $\Delta[BR]_{lim}$ remains 1.0 or close to 1.0 for $\frac{k_{RL}}{v_0} \geq 4$.

The solid red lines in Figure S2 represent fits to the different data sets using the function

(S18) $$\Delta[BR]_{lim} = A - \frac{1-A}{1-\exp\left(ln\frac{l_0}{l_{ref}}-12.5\right)}$$

where $l_{ref}$ is the reference rate constant (equal to 1.0) and *A* as a parameter that approaches 1.0 with increasing $\frac{k_{RL}}{v_0}$.

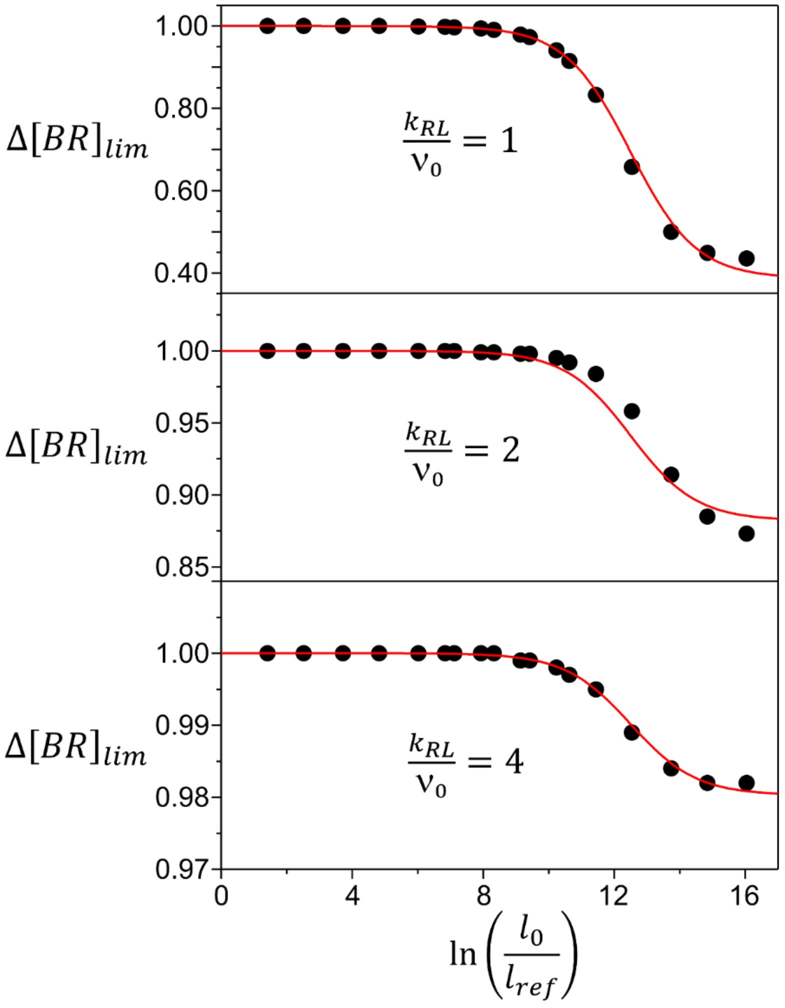


**Figure S2.** Limiting concentrations of BR after repetitive rotations of the cell as a function of the logarithm of $l_0$ for different $\frac{k_{RL}}{v_0}$ ratios.

**Table S1.** Relative concentrations of BR after passing through the laser beam Δ[BR] and the limiting value $\Delta[BR]_{lim}$ after repetitive rotations of the cell.[a]

| $P_0$/mW | $r_R$ /µm | $\frac{P_0}{r_R^2}$ /(mW/µm$^2$) | $l_0$/s$^{-1}$ | $\nu_0 = 5$ s$^{-1}$ | | | | $\nu_0 = 10$ s$^{-1}$ | | | | $\nu_0 = 25$ s$^{-1}$ | | | | $\nu_0 = 50$ s$^{-1}$ | | | |
|---|---|---|---|---|---|---|---|---|---|---|---|---|---|---|---|---|---|---|---|
| | | | | $\Delta t$/µs | Δ[BR] | $\Delta[BR]_{lim}$ $\frac{k_{RL}}{\nu_0}$ | | $\Delta t$/µs | Δ[BR] | $\Delta[BR]_{lim}$ $\frac{k_{RL}}{\nu_0}$ | | $\Delta t$/µs | Δ[BR] | $\Delta[BR]_{lim}$ $\frac{k_{RL}}{\nu_0}$ | | $\Delta t$/µs | Δ[BR] | $\Delta[BR]_{lim}$ $\frac{k_{RL}}{\nu_0}$ | |
| | | | | | | 4 | 2 | | | 4 | 2 | | | 4 | 2 | | | 4 | 2 |
| 2.5 | 2 | 2.5 | $9.280{\cdot}10^{6}$ | 9.794 | 0.030 | 0.983 | 0.858 | 4.897 | 0.094 | 0.984 | 0.870 | 1.959 | 0.187 | 0.986 | 0.880 | 0.979 | 0.236 | 0.986 | 0.885 |
| 0.75 | 2 | 0.75 | $2.784{\cdot}10^{6}$ | 9.794 | 0.053 | 0.984 | 0.866 | 4.897 | 0.146 | 0.986 | 0.881 | 1.959 | 0.265 | 0.988 | 0.896 | 0.979 | 0.333 | 0.989 | 0.910 |
| 0.25 | 2 | 0.25 | $9.280{\cdot}10^{5}$ | 9.794 | 0.140 | 0.987 | 0.889 | 4.897 | 0.290 | 0.989 | 0.909 | 1.959 | 0.451 | 0.992 | 0.932 | 0.979 | 0.566 | 0.994 | 0.953 |
| $7.5{\cdot}10^{-2}$ | 2 | $7.5{\cdot}10^{-2}$ | $2.784{\cdot}10^{5}$ | 9.794 | 0.413 | 0.992 | 0.935 | 4.897 | 0.586 | 0.995 | 0.954 | 1.959 | 0.733 | 0.997 | 0.972 | 0.979 | 0.820 | 0.998 | 0.983 |
| $2.5{\cdot}10^{-2}$ | 2 | $2.5{\cdot}10^{-2}$ | $9.280{\cdot}10^{4}$ | 9.794 | 0.712 | 0.997 | 0.972 | 4.897 | 0.818 | 0.998 | 0.981 | 1.959 | 0.895 | 0.999 | 0.990 | 0.979 | 0.961 | 0.999 | 0.994 |
| 10 | 30 | $1.1{\cdot}10^{-2}$ | $4.125{\cdot}10^{4}$ | 146.9 | 0.140 | 0.984 | 0.865 | 73.45 | 0.369 | 0.988 | 0.901 | 29.38 | 0.660 | 0.994 | 0.947 | 14.69 | 0.801 | 0.996 | 0.969 |
| $7.5{\cdot}10^{-3}$ | 2 | $7.5{\cdot}10^{-3}$ | $2.784{\cdot}10^{4}$ | 9.794 | 0.898 | 0.999 | 0.990 | 4.897 | 0.939 | 0.999 | 0.994 | 1.959 | 0.966 | 1.000 | 0.997 | 0.979 | 0.979 | 1.000 | 0.988 |
| 3 | 30 | $3.3{\cdot}10^{-3}$ | $1.237{\cdot}10^{4}$ | 146.9 | 0.545 | 0.992 | 0.929 | 73.45 | 0.735 | 0.995 | 0.959 | 29.38 | 0.880 | 0.998 | 0.981 | 14.69 | 0.934 | 0.999 | 0.990 |
| $2.5{\cdot}10^{-3}$ | 2 | $2.5{\cdot}10^{-3}$ | $9.280{\cdot}10^{3}$ | 9.794 | 0.964 | 1.000 | 0.997 | 4.897 | 0.979 | 1.000 | 0.998 | 1.959 | 0.989 | 1.000 | 0.999 | 0.979 | 0.993 | 1.000 | 0.999 |
| 1 | 30 | $1.1{\cdot}10^{-3}$ | $4.125{\cdot}10^{3}$ | 146.9 | 0.815 | 0.997 | 0.971 | 73.45 | 0.902 | 0.998 | 0.985 | 29.38 | 0.958 | 0.999 | 0.993 | 14.69 | 0.977 | 1.000 | 0.996 |
| $7.5{\cdot}10^{-4}$ | 2 | $7.5{\cdot}10^{-4}$ | $2.784{\cdot}10^{3}$ | 9.794 | 0.989 | 1.000 | 0.999 | 4.897 | 0.994 | 1.000 | 0.999 | 1.959 | 0.997 | 1.000 | 1.000 | 0.979 | 0.998 | 1.000 | 1.000 |
| 0.3 | 30 | $3.3{\cdot}10^{-4}$ | $1.237{\cdot}10^{3}$ | 146.9 | 0.940 | 0.999 | 0.991 | 73.45 | 0.969 | 0.999 | 0.995 | 29.38 | 0.987 | 1.000 | 0.998 | 14.69 | 0.993 | 1.000 | 0.999 |
| $2.5{\cdot}10^{-4}$ | 2 | $2.5{\cdot}10^{-4}$ | $9.280{\cdot}10^{2}$ | 9.794 | 0.996 | 1.000 | 1.000 | 4.897 | 0.998 | 1.000 | 1.000 | 1.959 | 0.999 | 1.000 | 1.000 | 0.979 | 0.999 | 1.000 | 1.000 |
| 0.1 | 30 | $1.1{\cdot}10^{-4}$ | $4.125{\cdot}10^{2}$ | 146.9 | 0.980 | 1.000 | 0.997 | 73.45 | 0.990 | 1.000 | 0.998 | 29.38 | 0.996 | 1.000 | 0.999 | 14.69 | 0.998 | 1.000 | 1.000 |
| $3.0{\cdot}10^{-2}$ | 30 | $3.3{\cdot}10^{-5}$ | $1.237{\cdot}10^{2}$ | 146.9 | 0.994 | 1.000 | 0.999 | 73.45 | 0.997 | 1.000 | 1.000 | 29.38 | 0.999 | 1.000 | 1.000 | 14.69 | 0.999 | 1.000 | 1.000 |
| $1.0{\cdot}10^{-2}$ | 30 | $1.1{\cdot}10^{-5}$ | 41.25 | 146.9 | 0.998 | 1.000 | 1.000 | 73.45 | 0.999 | 1.000 | 1.000 | 29.38 | 1.000 | 1.000 | 1.000 | 14.69 | 1.000 | 1.000 | 1.000 |
| $3.0{\cdot}10^{-3}$ | 30 | $3.3{\cdot}10^{-6}$ | 12.37 | 146.9 | 0.999 | 1.000 | 1.000 | 73.45 | 1.000 | 1.000 | 1.000 | 29.38 | 1.000 | 1.000 | 1.000 | 14.69 | 1.000 | 1.000 | 1.000 |
| $1.0{\cdot}10^{-3}$ | 30 | $1.1{\cdot}10^{-6}$ | 4.125 | 146.9 | 1.000 | 1.000 | 1.000 | 73.45 | 1.000 | 1.000 | 1.000 | 29.38 | 1.000 | 1.000 | 1.000 | 14.69 | 1.000 | 1.000 | 1.000 |

[a] Data was calculated for different rotational frequencies, $k_0/\nu_0$ ratios, and laser powers $P_0$ as described in the text.

### 2. Pump-Probe experiments

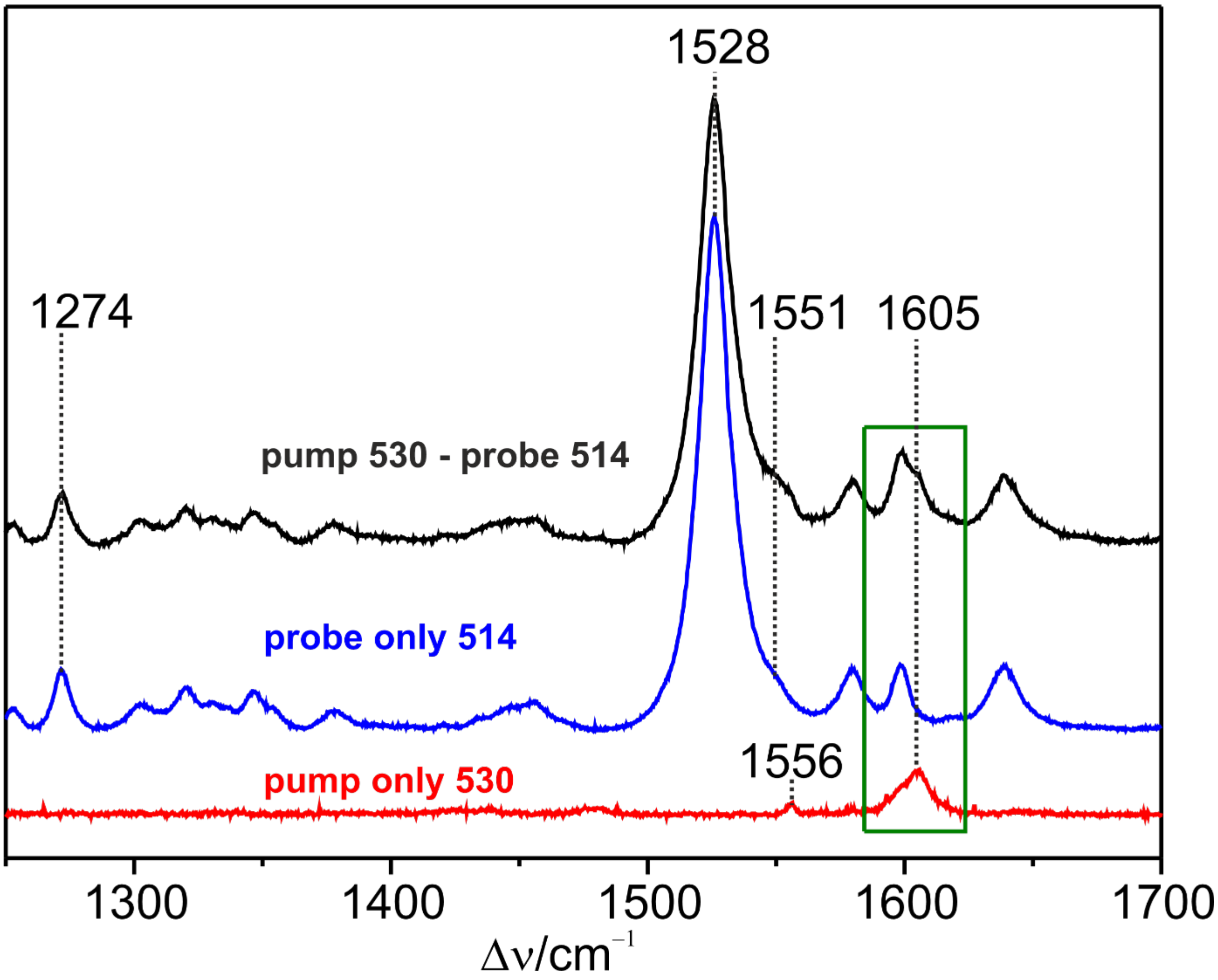


**Figure S3.** TR RR spectra of BR measured with the confocal set-up and a rotating cell frequency of 20 $s^{-1}$. The probe and pump wavelengths were 514.5 nm (0.1 mW) and 530.9 nm (100 mW), respectively. All three traces refer to same spectral region with respect to the 514.5 nm line (1250 – 1700 $cm^{-1}$). The black trace represents the pump-probe spectrum, including a small fraction of L550 as indicated by the enhanced intensity at 1551 $cm^{-1}$ compared to the probe only spectrum (blue trace). In addition, the pump-probe spectrum shows two bands at 1556 and 1605 $cm^{-1}$ which cannot be attributed to an intermediate. Instead, they correspond to the bands at the same position in the pump-only spectrum (red trace). These band results from the RR spectrum generated by the pump beam (530.9 nm – 18835 $cm^{-1}$), and measured on the scale of the probe beam (514.5 nm – 19436 $cm^{-1}$). Their frequencies agree very well with the 959 and 1009 $cm^{-1}$ bands of BR570, according to (19436 – 18836 + 956) $cm^{-1}$ = 1556 $cm^{-1}$ and (19436 – 18836 + 1009) $cm^{-1}$ = 1609 $cm^{-1}$.

### 3. Photocycles of Bacteriorhodopsin and Halorhodopsin

BR and HR are structurally very similar.[4] Kinetics and spectral properties compare rather well as far as the early reaction steps are concerned, which are in the focus of the present study (Fig. S4). Both reaction cycles depend on the pH.[4,6,7] Here, we have indicated approximate kinetic data for pH 7.0. In addition, the kinetic properties of HR also vary between the different organisms and strongly depend on the type of the anion and its concentration in the solution.[7,8] Here we have used HR from *Natronomonas pharaonis* and a chloride containing solution.

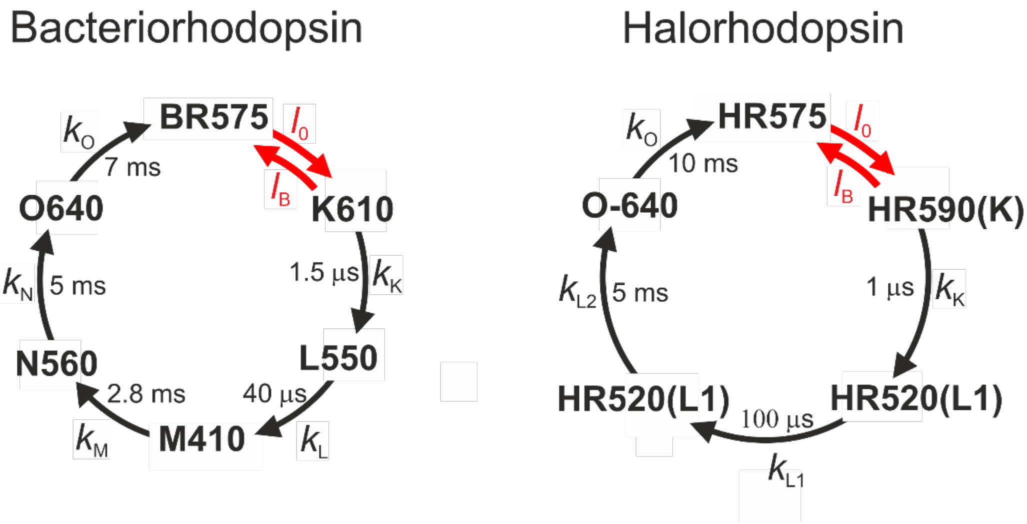


**Figure S4.** A, Simplified schemes of the photocycles of BR (left; see Fig. 1 in the MS) and HR from *Natronomonas pharaonis* (right). Spectral and kinetic data was taken from the literature.[4,6–8] The individual states are indicated by the commonly used notation with the numbers indicating the absorption maxima. The photochemical and thermal reactions are characterized by the red and black arrows/symbols, respectively. Back and cross reactions are neglected except for the primary photochemical process. The approximate life times of the individual reaction steps are indicated.
